# Unlocking inaccessible historical genomes preserved in formalin

**DOI:** 10.1101/2021.04.18.440380

**Authors:** Erin E. Hahn, Marina R. Alexander, Alicia Grealy, Jiri Stiller, Donald M. Gardiner, Clare E. Holleley

## Abstract

**Background:** Museum specimens represent an unparalleled record of historical genomic data. However, the wide-spread practice of formalin preservation has thus far impeded genomic analysis of a large proportion of specimens. Limited DNA sequencing from formalin-preserved specimens has yielded low genomic coverage with unpredictable success. We set out to refine sample processing methods and to identify specimen characteristics predictive of sequencing success. With a set of taxonomically diverse specimens collected between 1936 and 2015 and ranging in preservation quality, we compared the efficacy of several end-to-end whole genome sequencing workflows alongside a k-mer-based trimming-free read alignment approach to maximize mapping of endogenous sequence.

**Results:** We recovered complete mitochondrial genomes and up to 3X nuclear genome coverage from formalin-fixed tissues. Hot alkaline lysis coupled with phenol-chloroform extraction out- performed proteinase K digestion in recovering DNA, while library preparation method had little impact on sequencing success. The strongest predictor of DNA yield was overall specimen condition, which additively interacts with preservation conditions to accelerate DNA degradation.

**Conclusions:** We demonstrate a significant advance in capability beyond limited recovery of a small number of loci via PCR or target-capture sequencing. To facilitate strategic selection of suitable specimens for genomic sequencing, we present a decision-making framework that utilizes independent and non-destructive assessment criteria. Sequencing of formalin-fixed specimens will contribute to a greater understanding of temporal trends in genetic adaptation, including those associated with a changing climate. Our work enhances the value of museum collections worldwide by unlocking genomes of specimens that have been disregarded as a valid molecular resource.

## Background

Natural history collections are a window into the recent past, offering a view of historical biodiversity that is unparalleled in its detail. Collected over the last 250 years, voucher specimens document a period of time over which humans have had a devastating impact on the natural world (1). The comprehensive metadata associated with each specimen (collection date, location, sex, weight, age, etc.), phenotypic data (e.g., color, size, gut contents) and genomic data can be used to monitor ecosystem health and study the mechanisms driving adaptation, evolution, speciation and extinction (2, 3). The value of collections as sources of historical genetic material has been recognized for the past 30 years, with numerous pathways emerging to retrieve high-quality DNA from challenging archival vertebrate tissues such as skins (4), feathers (5, 6), eggshells (7, 8) and toe pads (9).

DNA degradation associated with preservation method and aging has limited most genetic studies of museum specimens to interrogation of relatively few loci via PCR amplification, often targeting the high copy mitochondrial genome. For phylogenetic studies where a survey of many-fold more loci improves understanding of species’ evolutionary history (10–12), genome-wide analyses are increasingly becoming common place. With demand for historical genome-wide data on the rise, newly-developed target-capture approaches now facilitate broader genomic survey from degraded museum specimens (13–15). In some cases, recovery and assembly of whole historical genomes has been achieved (16, 17), including, extinct from species (e.g., the Tasmanian tiger (18)). While technological advances are enabling recovery of genomic data from many museum specimens, genomic study of those preserved with 10% formalin (3.4% w/v formaldehyde) has thus far been very limited.

Formalin-fixation, followed by storage in ethanol, is a common curatorial method used to preserve soft tissue structure. Of the 1.9 million records of preserved chordates within the open-access Atlas of Living Australia (ALA) specimen database (19), 33% are classified as “spirit-preserved” (preserved in ethanol with or without prior formalin-fixation). A search for “formalin” preparation within the ALA’s chordate records indicates at least 4% of specimens (N = 77,301) have been formalin-fixed. This is likely a severe underestimate because formalin-fixation is not consistently recorded by all collections. Notably, for fish, reptiles and amphibians, formalin-fixation has historically been the primary method used to preserve tissues long-term while mammals and birds are commonly dry-preserved. Most collections now archive frozen fresh tissue specifically as a genomic resource. However, prior to the 1980s, spirit-preservation was the only method used to preserve soft tissue. Thus, spirit collections offer the only opportunity to obtain genetic data from a large proportion of older specimens, holotypes and some of the world’s most biodiverse vertebrate taxonomic groups.

Genomic study of formalin-preserved museum specimens has lagged behind because DNA extracted from such tissues is typically low-yield and highly fragmented. PCR amplification of formalin-degraded DNA templates is generally restricted to few, short genomic loci, which provide limited phylogenetic resolution (20). Formalin fixation presents further challenges by inducing numerous molecular lesions, such as strand breaks, base misincorporation, and both intra- and intermolecular cross-links (21–23). Formaldehyde damage to DNA templates can result in sequencing artefacts that are difficult to differentiate from true genetic variants (22, 23). Because PCR amplification of damaged DNA is particularly prone to sequencing artefacts, it is preferable to perform deep next-generation sequencing of amplicons (20) or to avoid amplicon approaches altogether through whole genome sequencing (WGS) of degraded templates (24). Coupled with library preparation methods optimized for low-input and damaged DNA templates (25, 26), high-throughput sequencing can generate enough coverage to call genomic variants with high confidence (27). Thus, WGS and reduced representations of genomes could provide a way to overcome the challenges associated with formalin damage and accurately reconstruct historical genetic variation from formalin-preserved tissues.

Promisingly, WGS of formalin-fixed paraffin-embedded (FFPE) archival tissues has become routine in clinical and medical contexts (28). However, museum specimens are often older, exposed to higher concentrations of formaldehyde, incubated in the fixative for longer (29) and in most cases have not been preserved in ideal conditions. Common museum practices, such as failure to rinse specimens prior to permanent storage in ethanol, result in prolonged formaldehyde exposure (30). Indeed, many specimens can be in contact with formaldehyde (or its derivatives, such as formic acid) for the entirety of their tenure in a collection. Prolonged formaldehyde exposure, especially under acidic conditions, is thought to result in more extreme DNA degradation (20, 31). The damage resulting from the preservation process compounds with DNA damage due to natural decomposition, which can be extensive and often precedes any obvious visual indicators of decomposition (32). Unfortunately, the time between death and preservation (post-mortem interval) is highly variable and rarely recorded. In light of these additional challenges, WGS methods used with FFPE tissues are relevant but not directly transferable to formalin-fixed museum tissues.

Of the few genetic studies of formalin-fixed museum specimens, most have targeted nuclear (33–37) and high copy mitochondrial (20,38,39) loci via PCR amplification due to the difficulty and unpredictability of nuclear DNA extraction. There are few examples of broader-scale genomic sequencing of formalin-fixed museum specimens and none have recovered whole vertebrate genomes. Hot alkaline extraction followed by WGS of a single 30-year-old formalin-preserved *Anolis* lizard yielded sufficient coverage to reconstruct the entire mitochondrial genome (40). Using the same method, whole genomes were recovered for the bioluminescent bacterial symbionts contained within light organs of formalin-preserved cardinalfish (41). Using a proteinase K digestion method, sufficient gDNA was recovered for capture and sequencing of ultra-conserved elements from formalin-preserved snakes (42). Hybridization capture baits have also been used to recover the mitochondrial genome from a 120-year-old formalin-preserved Crimean green lizard (43). Highlighting the difficulty of recovering gDNA from formalin-preserved specimens, numerous studies have reported failure to extract and amplify gDNA from formalin-preserved museum tissues (20,44,45). In this context, it is unfortunate yet wise to be hesitant to conduct destructive sampling of formalin-preserved specimens for the purposes of costly WGS.

Recent reports of successful, albeit limited, genomic sequencing in formalin-preserved specimens indicate WGS of higher quality specimens is possible. However, without a framework to guide specimen selection, genomic work on formalin-preserved museum tissues will remain infeasible. It is likely impossible to fully know the numerous and interdependent factors driving sequencing success (e.g., age of the specimen (46, 47), method of preservation (48), post-mortem interval (32) and heat and light exposure during storage). However, identification of metrics with which to pre-screen specimens for sequencing suitability will improve yield of genomic data while reducing unnecessary destruction of specimens. With screening criteria in hand, museum curators will be less reluctant to grant destructive sampling (49) and researchers will be more inclined to include historical specimens in their analyses.

To facilitate informed-selection of formalin-preserved museum specimens for WGS, we set out to further refine appropriate extraction and library preparation methods and to identify specimen characteristics predictive of DNA extraction and sequencing success. First, we investigated the relationship between residual formaldehyde concentration and pH in preservation media through a survey of specimens in the Australian National Wildlife Collection (ANWC; Crace, Australia). Next, in a phased approach, we compared DNA yield achieved with three extraction methods - (1) hot alkaline lysis digestion followed by phenol-chloroform extraction, (2) proteinase K digestion followed by phenol-chloroform extraction and (3) proteinase K digestion followed by silica spin column purification. We then applied the best-performing DNA extraction method to terrestrial vertebrate specimens representing the broad range of tissue quality observed in museum specimens and tested performance of two library preparation methods – (1) single-stranded method v2.0 (ss2) (25) and (2) BEST double-stranded method (dsBEST) (26). Placing our results into context with a comprehensive and unbiased survey of collection-wide spirit preservation conditions, we present a decision-making framework to accelerate and facilitate genomic research using formalin-preserved specimens.

## Results

### Preservation media condition survey

Within 149 ANWC specimen jars surveyed (23 amphibian, 40 mammal, 40 reptile, and 46 avian), preservation media pH ranged from 4.8–8.4 with 70 (47%), 61 (41%) and 18 (12%) having neutral (6.5–7.5), low (< 6.5) and high (> 7.5) pH, respectively. Residual formaldehyde concentration ([F]) ranged from 0–40,000 mg/L. High [F] (> 1000 mg/L) was detected in 61% of low pH jars, 6% of neutral pH jars and 0% of high pH jars. We assumed specimens in jars yielding [F] = 0 (n = 82) were preserved with ethanol and without exposure to formaldehyde. Consistent with the practice of fixing specimens with unbuffered formalin combined with the gradual degeneration of formaldehyde to formic acid, the pH of the formalin-preserved samples (range 4.8–7.1; mean = 6.2) was significantly lower than for the ethanol-preserved samples (range 6.1–8.4; mean = 7.1) (T-test; p < 0.0001; Supplementary Figure 1A). The recorded collection date of the specimens ranged from 1936–2015. The time since collection (age) of the ethanol-preserved specimens (mean = 40.1 years) was not significantly different than the formalin-preserved specimens (mean = 36.1 years) (Supplementary Figure 1B). Among the formalin-preserved samples, [F] and pH were negatively correlated (R = -0.6, p < 0.001; Figure 1). Age was not significantly correlated with either [F] or pH. Of the 12 specimens selected for sequencing, collection date ranged from 1962–2006 and pH ranged from 4.9–8.2. Three sequenced specimens were ethanol-preserved and nine sequenced specimens were formalin-preserved with [F] ranging from 325–20,000 mg/L (Table 1).

**Figure 1.**
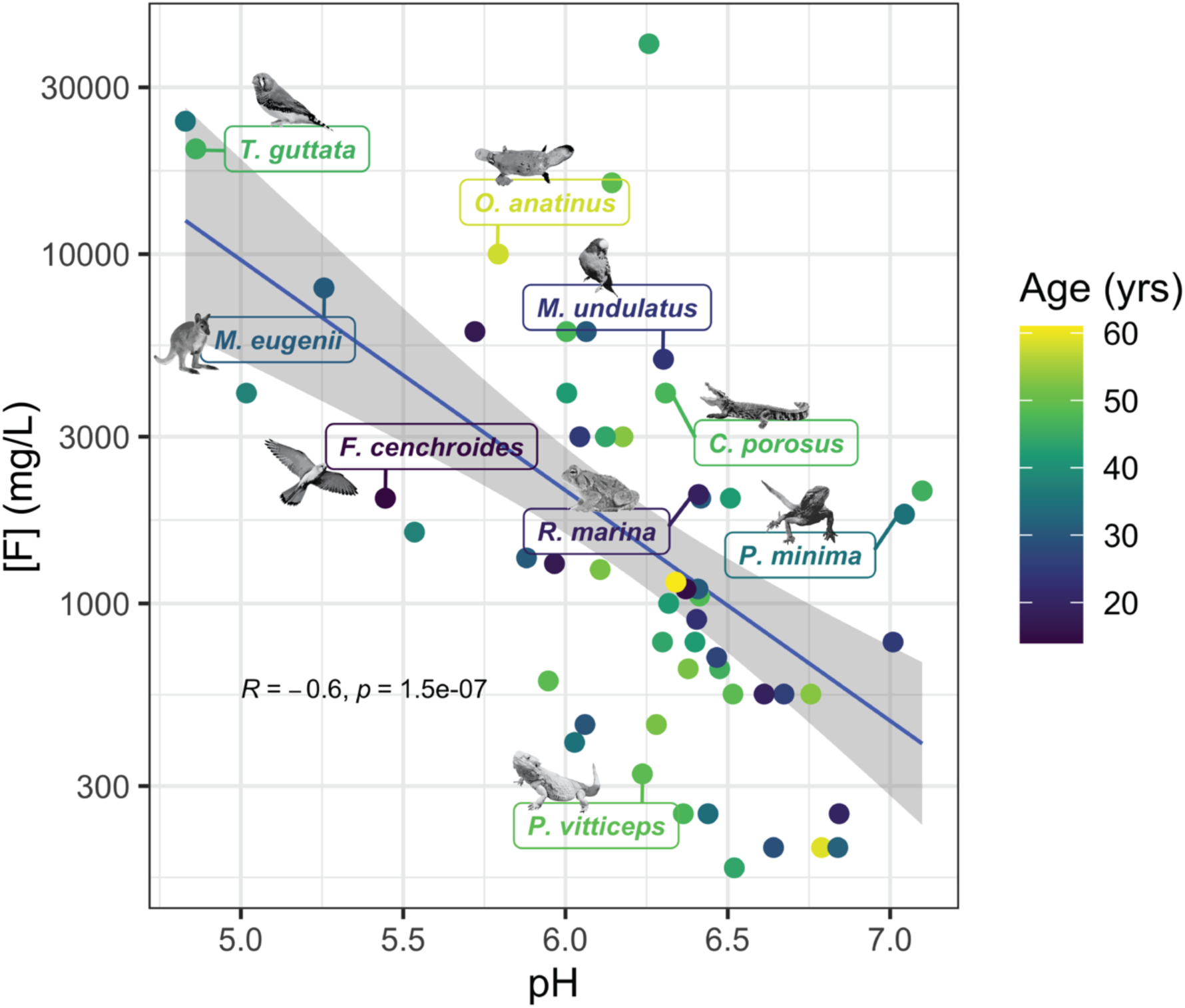
Preservation media survey results of formalin-fixed specimens in the Australian National Wildlife Collection. Residual formaldehyde concentration [F] (mg/L) is shown on a log-scale in relation to pH. Individual specimens (N = 65) are colored by the time since their collection (age) and the specimens selected for sequencing are indicated by species name. A linear model was used to fit a regression line and standard error is shown in grey; R = Pearson’s correlation coefficient.

**Table 1.**
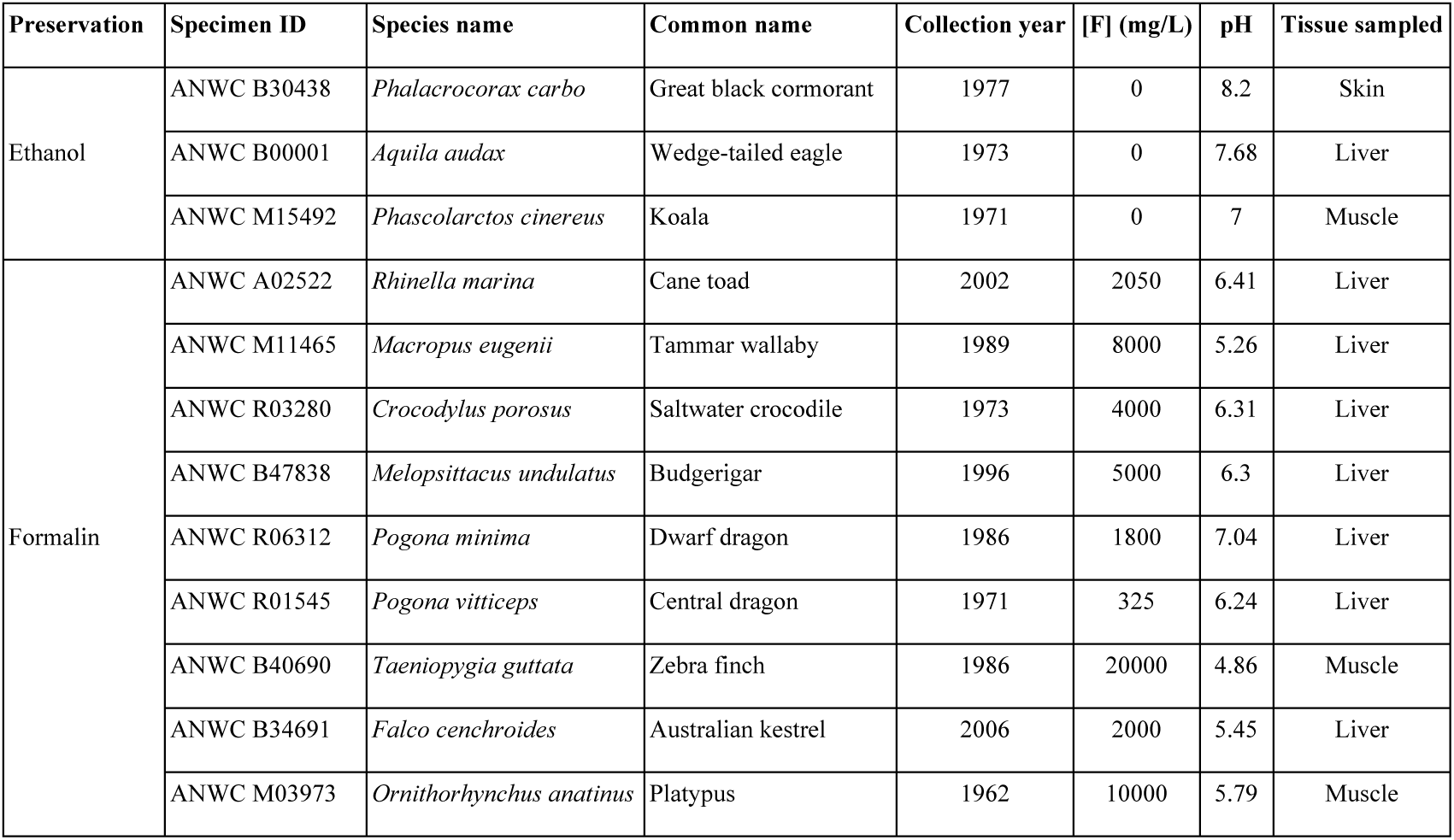
Specimen metadata and independently assessed preservation quality metrics for samples selected for sequencing. Twelve specimens (three ethanol-preserved and nine formalin-preserved) from the ANWC spirit vault were selected for DNA extraction and sequencing. Unique ANWC specimen IDs, species names, common name, recorded year of collection, residual formaldehyde concentration in the preservation media (mg/L), pH and tissue sampled for extraction are given.

### DNA quantification

We compared DNA yield from the hot alkaline lysis (HA), proteinase K plus phenol-chloroform (proK-PC) and proteinase K plus column (proK-col) extraction methods for the *Rhinella marina*, *Macropus eugenii* and *Crocodylus porosus* specimens and observed no significant differences between extraction methods (one-way ANOVA; Supplemental Figure 3A). However, the HA method produced more DNA from the two poor quality specimens (*M. eugenii* and *C. porosus*) compared to either of the proteinase K methods (Table 2). Thus, we predicted the HA method would perform better on specimens ranging broadly in preservation quality and we used this method to extract the remaining nine specimens. HA extraction yielded DNA detectable by high sensitivity Qubit for all twelve specimens. Two ethanol-preserved specimens (*Aquila audax* and *Phascolarctos cinereus*) and two formalin- preserved specimens (*R. marina* and *Melopsittacus undulatus*) yielded > 1,000 ng total DNA from 50 mg of tissue (Table 2). Three specimens, *Phalacrocorax carbo*, *Taeniopygia guttata* and *Ornithorhynchus anatinus*, yielded particularly low (< 100 ng) total DNA from 50 mg of tissue (Table 2). We observed no significant difference in DNA yield between ethanol and formalin-preserved specimens (T-test; Supplemental Figure 3B). However, mean DNA yield from ethanol-preserved specimens was more than double that from formalin-preserved specimens. Mean DNA yield from formalin-preserved specimens in preservation media with low pH (< 6) was not significantly different from those in media with neutral to high pH (> 6) (Supplemental Figure 3C). DNA yield was significantly higher from formalin-preserved liver tissue compared to non-liver tissue (T-test; p < 0.05; Supplemental Figure 3D). Both [F] and age showed a negative but non-significant correlation with DNA yield from formalin-preserved specimens (Supplemental Figures 3E and 3F).

**Figure 3.**
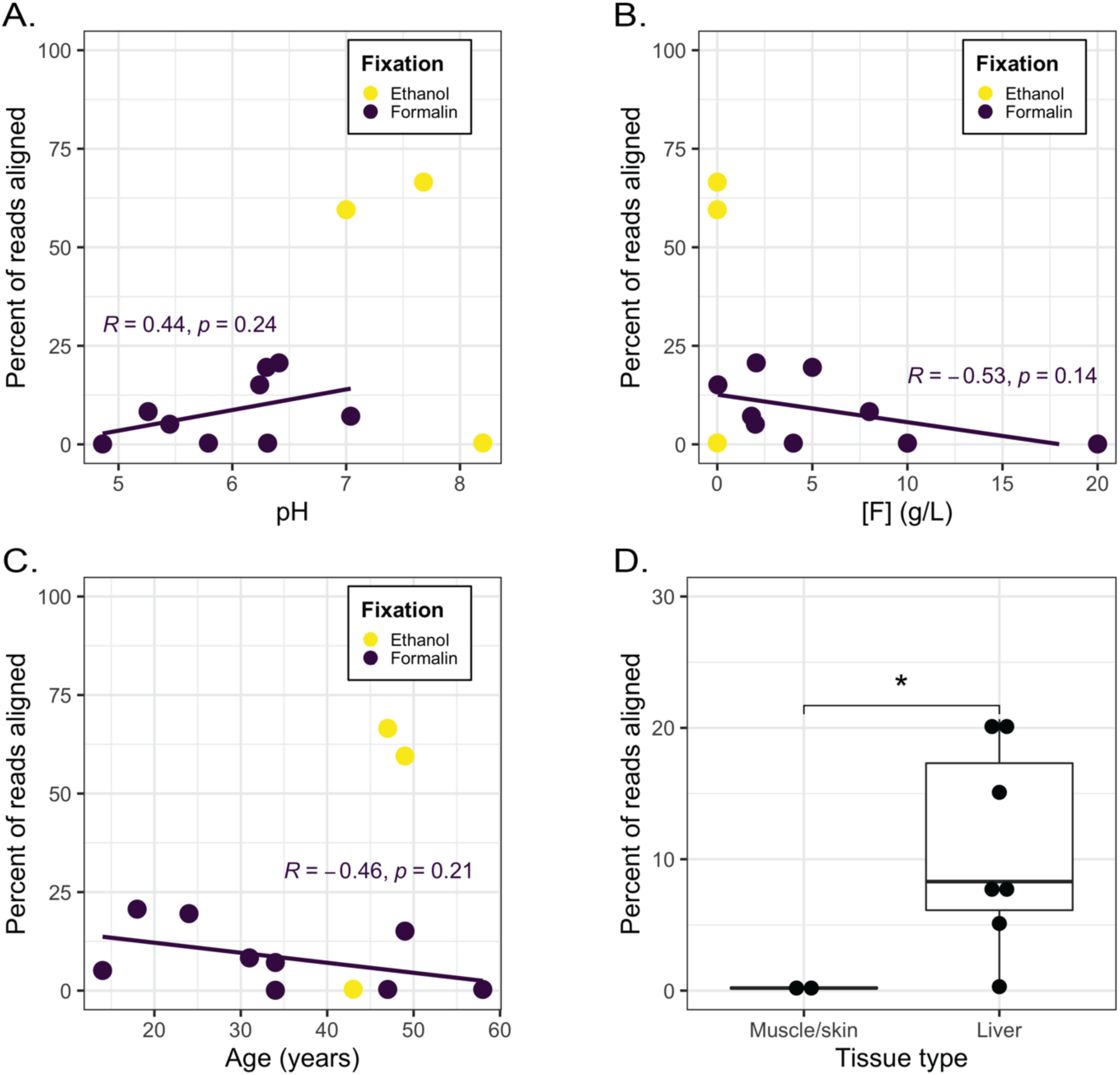
Alignment results for hot alkali extracted samples. The correlation between the percentage of reads aligned to the whole genome (combining both library preparations of the hot alkali extracted specimens) and (A) preservation media pH, (B) preservation media formaldehyde concentration (g/L), (C) number of years in the collection and (D) tissue sampled. In A-C, all specimens are shown colored by their fixation type and R = Pearson’s correlation coefficient for the formalin-fixed specimens. In D, only the formalin-preserved specimens are plotted and individual specimens are shown with black dots, * = p < 0.05.

**Table 2.** Sequencing and alignment statistics. For all specimens, DNA yield is given for the individual extractions of 50 mg of tissue. For the remaining metrics, the values shown were calculated having combined both the ss2 and dsBEST libraries. The number of raw reads is given as a sum of all single reads (R1 and R2) from the paired-end sequencing run. Reads aligned indicates the percent of raw reads aligned to reference genome after removal of PCR and optical duplicates. The mean aligned insert length is the mean length (in bp) of the aligned portion of the read. C_nuc_ is the coverage of the nuclear genome. C_mt_ is the proportion of mitochondrial genome with greater than 30X coverage. C_pot_ is the estimated potential genomic coverage if the full library had been sequenced, calculated from the estimated library complexity. MRM is the number of reads aligned to the mitochondrial genome per one million raw reads.

| Preservation | Species | Extraction method | DNA yield from 50 mg (ng) | Raw reads (million) | Reads aligned (%) | Mean aligned insert length (bp) | $C_{nuc}$ | $C_{mt}$ | $C_{pot}$ | MRM |
| --- | --- | --- | --- | --- | --- | --- | --- | --- | --- | --- |
| Formalin | *Rhinella marina* | HA | 1,860 | 434 | 21 | 65 | 2.2 | 0.78 | 6.2 | 52 |
|  |  | proK-col | 666 | 77 | 40 | 81 | 1 | 0 | 6.2 | 14 |
|  |  | proK-PC | 2,550 | 321 | 15 | 74 | 1.2 | 0.42 | 11.4 | 29 |
|  | *Macropus eugenii* | HA | 271 | 306 | 8 | 56 | 0.5 | 0.59 | 2.7 | 50 |
|  |  | proK-col | 4 | 17 | 1 | 67 | 0 | 0 | 0.1 | 3 |
|  |  | proK-PC | 33 | 801 | < 1 | 65 | 0 | 0 | 0.1 | 2 |
|  | *Crocodylus porosus* | HA | 130 | 23 | < 1 | 67 | 0 | 0 | 0 | 11 |
|  |  | proK-col | None detected | 160 | < 1 | 70 | 0 | 0 | 0.1 | 12 |
|  |  | proK-PC | 79 | 294 | < 1 | 62 | 0 | 0 | 0 | 2 |
|  | *Melopsittacus undulatus* | HA | 2,400 | 318 | 20 | 60 | 3.1 | 0.94 | 23.6 | 201 |
|  | *Pogona minima* | HA | 521 | 367 | 7 | 58 | 0.8 | 0.51 | 7.5 | 29 |
|  | *Pogona vitticeps* | HA | 672 | 432 | 15 | 59 | 2.1 | 0.85 | 7.9 | 52 |
|  | *Taeniopygia guttata* | HA | 15 | 62 | < 1 | 66 | 0 | 0 | 0 | 1 |
|  | *Falco cenchroides* | HA | 690 | 303 | 5 | 56 | 0.7 | 0.12 | 2.1 | 14 |
|  | *Ornithorhynchus anatinus* | HA | 22 | 520 | < 1 | 70 | 0 | 0.13 | 0.8 | 20 |
| Ethanol | *Phalacrocorax carbo* | HA | 57 | 292 | < 1 | 69 | 0.10 | 0.90 | 0.60 | 50.00 |
|  | *Aquila audax* | HA | 1,932 | 282 | 67 | 76 | 11.3 | 0.98 | 323 | 2515 |
|  | *Phascolarctos cinereus* | HA | 1,254 | 423 | 60 | 76 | 5.4 | 0.94 | 93 | 2606 |

### Pre-alignment library quality assessment

Prior to alignment, we used FastQC to assess the quality of paired-end reads from ss2 and dsBEST libraries. All libraries contained a high proportion of adapter content and low read quality score beginning at roughly 50 bp, consistent with highly fragmented input DNA. Focusing on the first 75 bp of the raw reads, mean sequence quality was slightly but significantly higher for read 2 (mean Phred score = 34.3) than for read 1 (mean Phred score = 33.7) across all libraries (paired T-test; p < 0.001). Likewise, the mean sequence quality was significantly higher in ss2 libraries compared to the corresponding dsBEST libraries for both read 1 (mean of the differences = 2.1; paired T-test; p < 0.001) and read 2 (mean of the differences = 0.79; paired T-test; p < 0.01). Mean sequence quality was not significantly different between reads derived from ethanol and formalin-preserved tissues, even when excluding libraries prepared from less than 200 ng of input DNA (paired T-test). We found evidence of cross-contamination in several libraries prepared from low DNA yield extractions. Compared to negative controls, both *O. anatinus* libraries and all but two *C. porosus* libraries showed a higher number of reads classified as genus *Mus* by Kraken2 (Supplementary Table 2). The *O. anatinus* libraries also contained a high percentage of reads classified as *Homo sapiens* (9.7% and 25%). The *O. anatinus* and *C. porosus* tissues were among those that yielded the least DNA. The *O. anatinus* HA extraction yielded just 22 ng. The *C. porosus* HA and proK-PC extractions yielded 130 and 79 ng, respectively, while the proK-col extraction yielded no detectable DNA. The only other specimens to yield less than 500 ng were the *P. carbo*, *T. guttata* and *M. eugenii*.

### Relative alignment quality from three extraction methods

We used three indicators of alignment quality to compare the relative success of the three extraction methods on the *R. marina*, *M. eugenii* and *C. porosus* specimens: percent of raw reads aligned to the genome (% alignment), the number of reads aligned to the mitochondrial genome per million raw reads (MRM) and the mean aligned insert length. Among these three specimens, we observed no significant differences between library preparation methods in any of the three alignment quality indicators (paired T-tests). Therefore, we took the mean of the two library preparations to compare extraction methods across each alignment quality indicator. Again, we observed no significant difference between the three extraction methods applied to the *R. marina*, *M. eugenii* and *C. porosus* specimens in any of the three alignment quality indicators (one-way ANOVA). All six *C. porosus* libraries yielded < 1% alignment (Figure 2A and Table 2), indicating failure of all extraction and library preparation methods on this specimen. Excluding the *C. porosus* libraries, we observed significant differences in MRM between the extraction methods (one-way ANOVA; p < 0.05) with the HA method producing significantly more MRM than both the proK-col and proK-PC methods (Tukey tests; p < 0.05). We observed no significant difference in MRM between the proK-col and proK-PC methods (Tukey tests) nor in % alignment or mean insert length between the three extraction methods (one-way ANOVA).

**Figure 2.**
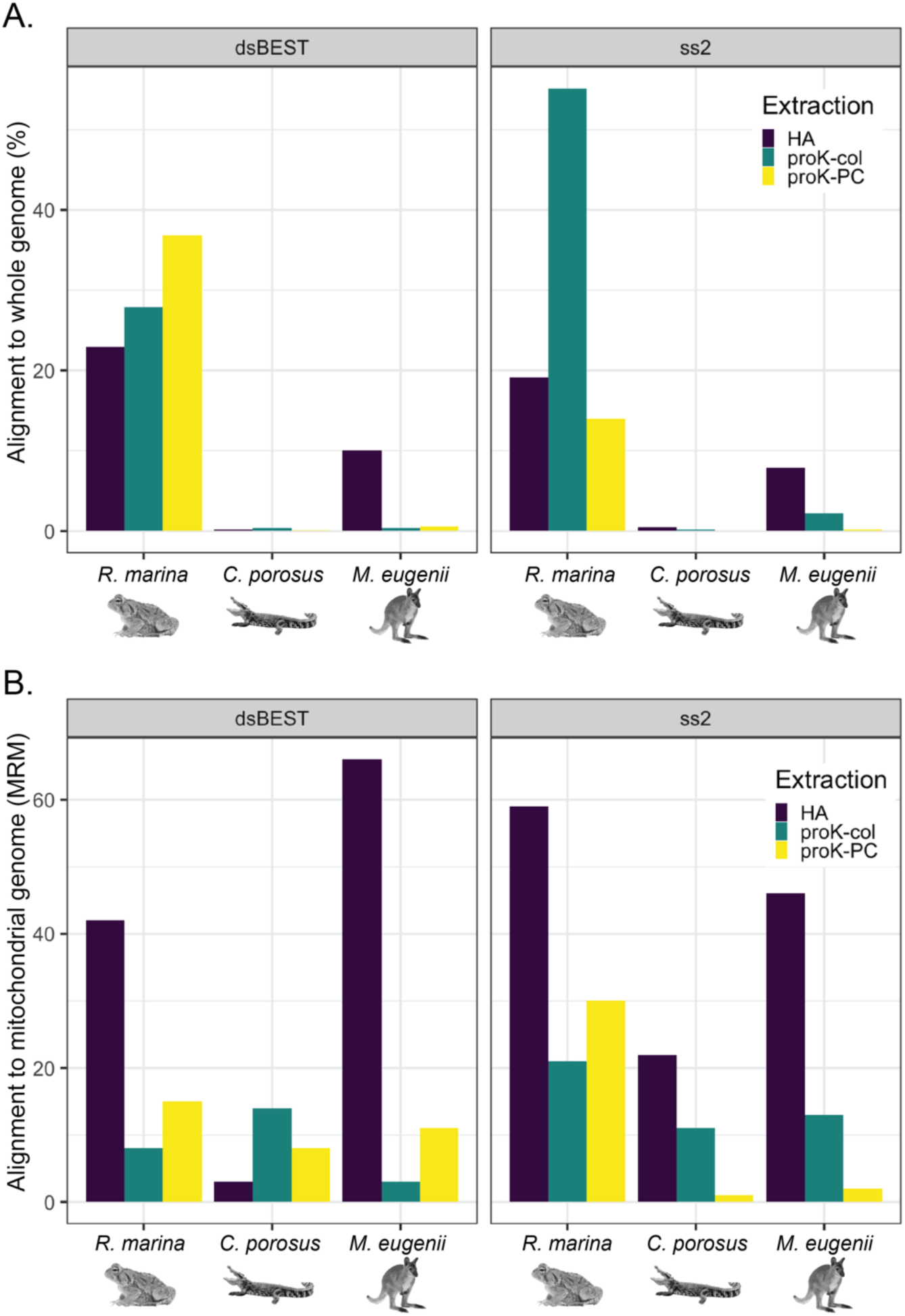
Effectiveness of extraction and library preparation methods for *R. marina*, *C. porosus* and *M. eugenii* specimens. (A) Alignment to the whole genome expressed as the percentage of reads aligning (B) Alignment to the mitochondrial genome expressed as the number of reads aligned per million raw reads (MRM). dsBEST = BEST double-stranded method (26); ss2 = single-stranded method v2.0 (25); HA = hot alkaline lysis; proK-col = proteinase K digestion followed by column purification; proK-PC = proteinase K digestion followed by phenol-chloroform extraction.

### Effect of specimen quality on sequencing success

The percentage of aligned reads removed by optical and PCR de-duplication varied between 8.8% and 99.5% across all libraries. Among the HA alignments, de-duplication reduced significantly more mapped reads from dsBEST libraries than from ss2 libraries (paired T-test; p < 0.01). Combining the ss2 and dsBEST libraries for each HA extraction, de-duplication removed more than double the percentage of reads (69.8% versus 32.8%) from poor quality specimens (those yielding < 1% reads aligned) compared to better quality specimens (those yielding > 1% reads aligned). However, this difference was not significant (T-test). De- duplication removed significantly more reads from the formalin-preserved specimens (mean = 54.6%) than from the ethanol-preserved specimens (mean = 16.7%) (T-test; p < 0.01). Following de-duplication, the mean percent of mapped reads remaining was 44% and 59% for the dsBEST and ss2 HA libraries, respectively. Across all specimens extracted using the HA method, we observed no significant differences between library preparation methods in any of the three alignment quality indicators (paired T-tests). Therefore, we conducted further comparison of the effect of specimen quality on alignment success taking the mean of each alignment quality indicator from the two HA library preps.

HA extraction of one of three ethanol-preserved specimens (*P. carbo*) and three of nine formalin-preserved specimens (*C. porosus*, *T. guttata* and *O. anatinus*) produced < 1% aligned reads (Table 2), indicating equal rates of very poor sequencing success with ethanol- and formalin-preserved tissues. Excluding the specimens with < 1% aligned reads, the ethanol-preserved specimens produced a significantly higher percentage of aligned reads (T-test; p < 0.01). Two of the three ethanol-preserved specimens (*A. audax* and *P. cinereus*) produced > 60% aligned reads while the remaining six formalin-preserved specimens (*R. marina*, *M. eugenii*, *M. undulatus*, *Pogona minima*, *Pogona vitticeps* and *Falco cenchroides*) produced between 5% and 21% aligned reads (Table 2). Excluding the specimens with < 1% aligned reads, the mean insert length was significantly longer for the ethanol-preserved specimens (mean = 76 bp) compared to the formalin-preserved specimens (mean = 59 bp) (T-test; p < 0.0001). MRM was also significantly higher for the ethanol-preserved specimens (mean = 2,560) compared to the formalin-preserved specimens (mean = 43) (T-test: p < 0.01).

The percentage of reads aligned increased with preservation media pH (R = 0.44; Figure 3A), decreased with preservation media [F] (R = -0.53; Figure 3B) and decreased with specimen age (R = -0.46; Figure 3C), although these correlations were not statistically significant. The percentage of aligned reads was significantly higher in specimens sampled with liver than those sampled with muscle and skin (T-test; p < 0.05; Figure 3D). Of the specimens yielding poor sequencing success (< 1% reads aligned), all but *C. porosus* were sampled with either muscle or skin as liver was not present. The only specimen sampled with a tissue other than liver to yield a percent of reads aligned > 1% was the ethanol-preserved *P. cinereus*.

### Genome sequencing coverage

Nuclear genome coverage (C_nuc_) of the deduplicated alignments was < 1X for the majority of libraries. Since raw read yield was highly variable, C_nuc_ is not an appropriate measure with which to compare the extraction or library preparation methods. However, it is noteworthy that we achieved C_nuc_ > 1X for two of the ethanol-preserved specimens and three of formalin-preserved specimens. Combining all libraries for a given specimen, we achieved a total of 5.4X and 11.3X C_nuc_ for the ethanol-preserved *P. cinereus* and *A. audax* specimens, respectively (Table 2). Likewise, we achieved a total of 2.1X, 3.1X and 4.4X C_nuc_ for the formalin-preserved *P. vitticeps*, *M. undulatus* and *R. marina* specimens, respectively (Table 2). To estimate the potential for improving C_nuc_ through re-sequencing of the prepared libraries, we calculated potential genomic coverage (C_pot_) (Table 2). Combining all libraries for a given specimen, C_pot_ exceeded 20X for the *R. marina* and *M. undulatus* and exceeded 75X for the *P. cinereus* and *A. audax*. Focussing on the mitochondrial genome, the proportion of sites with 30X or higher coverage (C_mt_) was nearly complete (> 0.9) for all three ethanol-preserved specimens (Table 2). C_mt_ for the formalin-preserved *M. undulatus* (0.94) was comparable to that of the ethanol-preserved specimens. C_mt_ was moderate to high (> 0.5) for five of the formalin-preserved specimens (Table 2). Only the *C. porosus*, *T. guttata*, *F. cenchroides* and *O. anatinus* yielded very poor C_mt_ (< 0.15).

### Read length periodicity

From the aligned insert lengths estimated with Picard, we plotted the frequency of reads between 50 and 100 bp (Figure 4). This plot revealed a pattern of read length periodicity in several specimens, notably those that resulted in higher mapping success. We observed prominent periodicity of approximately 10.1 bp in the *R. marina* specimen extracted with the proK-PC method. While less pronounced, we observed read length periodicity of approximately 10.8 bp in the HA extractions of *R. marina*, *P. vitticeps*, *P. minima*, *F. cenchroides*, *A. audax* and *P. cinereus*. The pattern of periodicity was observed in both the dsBEST and ss2 libraries, however, it was slightly more pronounced in the dsBEST libraries.

**Figure 4.**
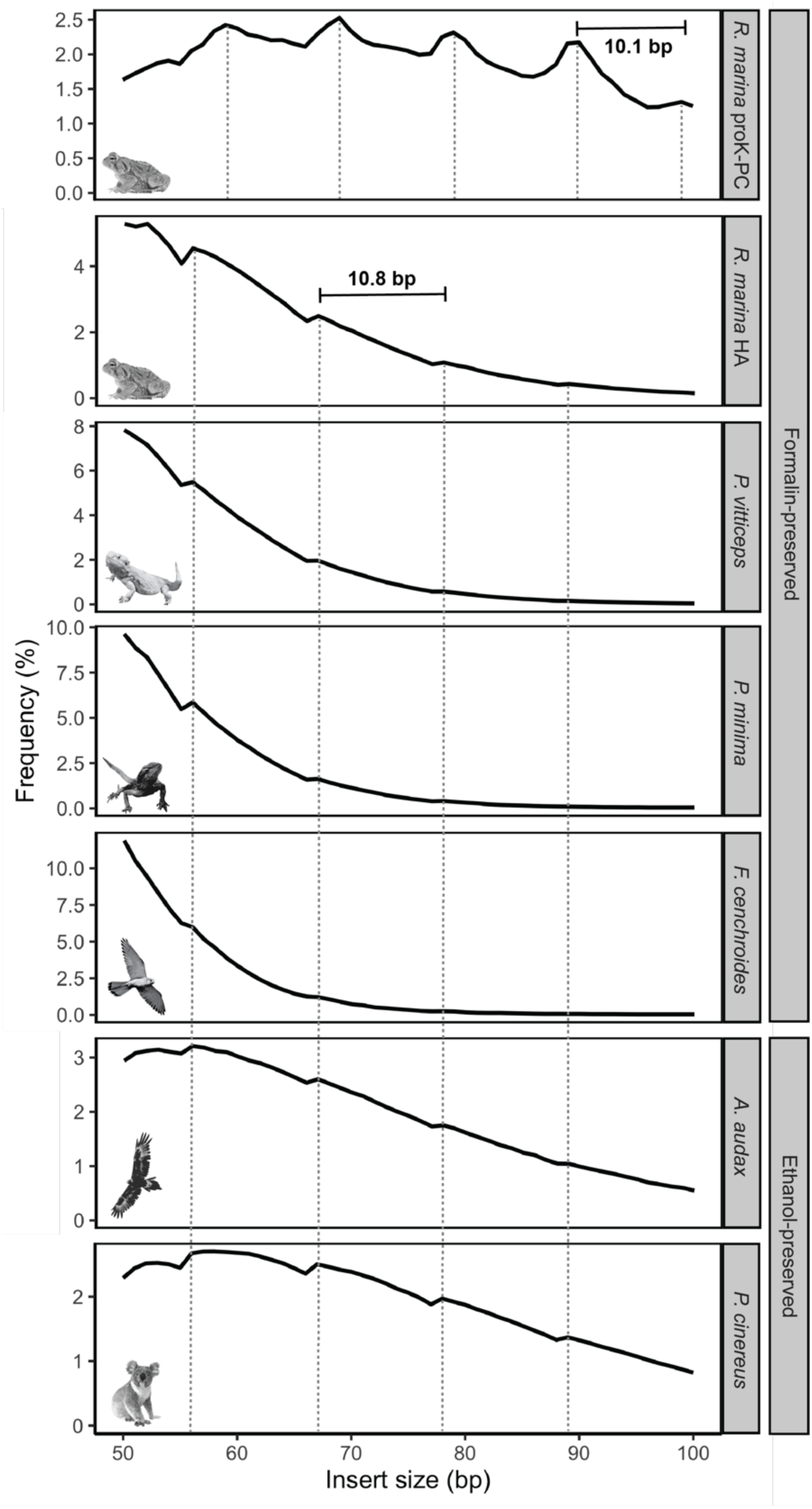
Libraries with read periodicity. The frequency of insert lengths, in bp, estimated from the mapped dsBEST libraries is shown for six preserved specimens. Read periodicity in the *R. marina* libraries from the proteinase K with phenol-chloroform (proK-PC) extractions averages 10.1 bp while periodicity in libraries from the hot alkali extractions of six specimens averages 10.8 bp.

## Discussion

In this study, we present evidence challenging the common perception that formalin-preserved museum specimens are devoid of accessible DNA. Processed with a tailored molecular and bioinformatic workflow, formalin-preserved specimens had an overall sequencing success rate equivalent to ethanol-preserved specimens, albeit with recovery of a lower percentage of sequence reads mapping to the reference genome. Contrary to popular belief, we found genome-wide nuclear data is retrievable from some formalin-preserved museum specimens, even with a moderate investment of sequencing effort (with 30% of formalin-preserved specimens, we achieved > 2X nuclear genome coverage from 300-500 million raw reads). We also show reconstruction of large sections of the mitochondrial genome is possible even in poor quality specimens where limited nuclear data were recovered (with 55% of formalin-preserved specimens, we achieved > 30X coverage of more than 50% of the mitochondrial genome). Investigating specimens covering a range of preservation quality, we also developed a decision-making framework to improve sequencing success rate and prioritize suitable specimens. Our findings support a considered and targeted sequencing approach that transforms thousands of spirit collection specimens into a new molecular resource. Improved access to genomic data held in these specimens has the potential to inform research into the mechanisms driving adaptation, evolution, speciation and extinction.

### Hot alkaline lysis effectively recovers gDNA from formalin-preserved archival tissues suitable for next generation sequencing

Originally developed for DNA extraction from FFPE sections, the HA method relies on high heat (120°C) under alkaline conditions (pH = 13) to break strong inter- and intramolecular cross links and utilizes organic extraction to maximize capture of fragmented gDNA from formalin-preserved tissues (50–52). This method has been applied to museum specimens to successfully recover sections of the mitochondrial genome in trout (53) and full mitochondrial genomes from lizards (40) and bacterial symbionts (41). Here we show the HA yields gDNA in adequate quantities for WGS from higher-quality formalin-preserved museum specimens. Coupled with library preparation methods designed to efficiently convert degraded DNA, we produced complex sequencing libraries with the potential to recover full vertebrate genomes when mapped using a strategy optimized to maximize recovery of endogenous sequence. Our results indicate that the HA method is appropriate for DNA extraction from a broad range of taxa preserved under various conditions, making it well-suited for application in both museum and pathological settings.

In a small-scale comparison to proK digestion with either phenol-chloroform extraction or column purification, the HA method performed superiorly for poor quality formalin-preserved specimens. We experienced equal success rates with the HA method in formalin and ethanol-preserved tissues. It is not standard practice to apply the HA method to ethanol-preserved specimens, which do not suffer from cross-linking, but we implemented it in this study to serve as a comparison to formalin-fixed tissues. Thus, while the HA method is likely unnecessarily harsh for recovery of DNA from tissues not crosslinked with formaldehyde, we propose this extraction method is suitable across a wide range of tissue qualities and preservation conditions observed in museum spirit collections. And, given that we achieved relatively high yield from the ethanol-preserved tissues, we propose that the HA method is appropriate in cases where contact with formalin cannot be determined. We caution; however, the HA method’s success may be limited to DNA-rich tissues such as liver. Our HA extractions of formalin-preserved muscle and ethanol-preserved skin tissue failed to yield adequate gDNA for sequencing, while our HA extraction of ethanol-preserved muscle tissue was less successful than our extraction of ethanol-preserved liver tissue. HA extraction has been previously observed to perform poorly compared to cetyltrimethylam-monium bromide (CTAB) protocols on formalin-preserved mammalian heart tissue (54). We also note that, preservation conditions being equal, DNA yield may differ between taxonomic groups due to factors such blood cell nucleation. Due to low sample size, we were not able to test if the lack of nucleated red blood cells in mammal tissues impacted DNA yield.

### aDNA library preparation methods effectively capture DNA extracted from formalin-preserved archival tissues

DNA degradation in museum specimens is a significant challenge to genome sequencing. To improve our conversion of degraded DNA from formalin-preserved tissues into high quality library molecules, we utilized two library preparation methods developed specifically for degraded aDNA templates. We tested the ss2 (46) and dsBEST (47) methods on DNA extracted from both ethanol and formalin-preserved archival tissues. Sequence quality was significantly higher for libraries prepared using the ss2 method compared to the dsBEST protocol. However, this quality difference did not result in significantly lower rates of read alignment or reduced mapped insert length for the dsBEST libraries. While we did not see differences in contamination rates between the two methods, an advantage of the dsBEST method is its reliance on fewer tube transfers and additions of solution, thus reducing opportunities to lose DNA and introduce contaminants. The ss2 and dsBEST methods performed similarly on all twelve of our archival templates, indicating both are well-suited to prepare libraries from DNA extracted from ethanol and formalin-preserved tissues. Alternative library preparation methods developed specifically for degraded DNA may prove equally effective. To maximize conversion of fragmented archival DNA template, we advise using a library preparation method designed to capture small fragments whilst minimising contamination risk. Overall, we observed samples with very low DNA yield (< 200 ng from 50 mg of tissue) did not produce libraries with high rates of mapping success. Thus, as a cost-saving measure, we advise quantifying DNA templates prior to library preparation and focussing sequencing effort on higher yielding samples.

### High alignment rates of fragmented DNA are achieved through exhaustive match searching

Removal of adapter sequence and low-quality bases via read-trimming is a standard pre- processing procedure conducted on raw sequencing reads prior to mapping. In the context of libraries prepared from highly degraded templates, filtering and trimming can reduce the dataset substantially. For example, pre-processing of the library prepared from a formalin-preserved *Anolis* lizard reduced the dataset to 13.5% of the raw data (40). Although filtering and trimming are effective at removing PCR duplicates and erroneous bases introduced through library preparation and sequencing, quality control parameters should be optimized to avoid removing informative endogenous sequence, particularly with data derived from highly fragmented low-input templates. Compared to DNA extractions from fresh tissue, our extractions from formalin-preserved specimens were highly fragmented as is typical of aDNA sources (55). We opted to trial a computationally efficient approach that eliminates loss of endogenous sequence during pre-processing. The kalign function from the open source kit4b toolkit performs alignments of raw reads by searching for the maximum length match within the read to the reference sequence regardless of the match’s position within the read. For each raw read, kalign performs a rapid complete exhaustive match search across the indexed reference genome. The match search is performed recursively through seed expansions generated along the read length. The longest match to endogenous sequence is retrieved while satisfying the minimum length threshold of the match. Using this approach, we aligned up to 21% and 67% of raw reads from formalin and ethanol-preserved tissues, respectively. These alignment rates are consistent with the degree of degradation in the DNA we extracted from spirit-preserved museum specimens being intermediate between that of fresh and truly ancient tissues. A previous application of the ss2 method yielded a maximum of 11.3% mappable reads from libraries prepared from aDNA tissue sources (25). The same study yielded 60% and 68% mappable reads from libraries prepared from horse and pig liver stored in buffered formalin for 5 and 11 years, respectively (25). In comparison, our modest alignment rates may be the result of tissues of intermediate age and using a different metric of calculating the percent of mapped reads.

### Sequencing success is strongly influenced by specimen integrity prior to fixation

To explore the effects of formalin-fixation on sequencing success, we selected three specimens preserved with ethanol only and nine specimens preserved with formalin. We found no significant difference in DNA yield between the ethanol and formalin-preserved specimens and the differences we observed in DNA fragment lengths were minimal. Furthermore, we observed equal rates of very poor sequencing success within ethanol and formalin-preserved specimens, indicating preservation method is not a strict determinant of sequencing success. Older, poor-quality ethanol-preserved specimens have previously been shown to be as problematic for genomic analyses as formalin-preserved specimens (42, 56). This is not to say preservation method does not impact sequencing success. Two of our ethanol-preserved specimens (*P. cinereus* and *A. audax*) had much higher mapping rates (60% and 67% reads aligned, respectively) than even our most successful formalin-preserved specimens (*R. marina,* produced 21% reads aligned with the HA method). Our findings indicate WGS of formalin-preserved museum specimens is possible using HA extraction paired with a library preparation optimized for conversion of degraded DNA. However, as with all potential DNA sources, the overall integrity of the tissue will ultimately determine sequencing success.

The specimens with poor sequencing success (< 1% reads aligned) were largely older, their preservation media had lower pH and higher [F] and they were sampled with a tissue other than liver. On the contrary, the specimens with better sequencing success were preserved more recently, their preservation media had neutral pH and lower [F] and the tissue sampled was liver. We calculated the correlation between specimen quality measures ([F], pH, age and tissue type) and both DNA yield and mapping success. Tissue type was the only quality measure significantly associated with lower DNA yield, with liver yielding significantly more DNA than either muscle or skin. Our higher success with liver is consistent with findings of a previous study comparing sequencing success from liver, muscle and tail-tip in a formalin-preserved *Anolis* lizard (40). However, in that study, the tissues were extracted using different methods and thus it could not be determined if success was driven by tissue type or extraction method.

Post-mortem DNA degradation occurs more rapidly in liver relative to other bodily tissues including skeletal muscle, heart and brain (57, 58). In the museum curatorial setting, specimens undergo varying degrees of post-mortem decay prior to fixation. As is the case for most museum specimens, the length of the post-mortem interval (PMI) was not recorded for the specimens used in this study. Given expected rapid decay of the viscera, we used the visual appearance of the gut contents as a reasonable proxy for the length of the PMI. The four specimens used in this study that lacked liver tissue were visibly more degraded than those with intact liver tissue (Supplementary Figure 2). In the case of the *P. cinereus*, *P. carbo* and *O. anatinus*, the complete absence of viscera indicated the internal organs were likely well- degraded and discarded prior to fixation. For specimens preserved after a long PMI, DNA integrity throughout the carcass would be lower than in specimens preserved after a short PMI.

Therefore, we conclude that the higher yield from specimens sampled with liver is a reflection of overall specimen quality and DNA damage occurring post-mortem but prior to fixation.

### Re-thinking formalin damage

Formalin-preserved museum specimens have long been considered intractable sources of gDNA. Encouragingly, we found specimen contact with formaldehyde does not prohibit DNA sequencing if tissue decomposition occurring prior to fixation is minimized. With appropriate sample vetting (Figure 5), HA extraction and DNA library preparation optimized for degraded DNA, historical genomic data may be extracted from many formalin-preserved specimens. These data will not be of similar quality to those recovered from fresh or ethanol-preserved tissues. However, higher sequencing volume and borrowing of analytical methods from the field of aDNA may facilitate reconstruction of historical genomes from formalin-preserved tissues. We found evidence that DNA damage in formalin-preserved specimens shares characteristics with that of aDNA. In addition to capturing shorter fragments with low mapping rates, we observed a pattern of read length periodicity of approximately 10 bp. This is consistent with observations in aDNA specimens (59) and is an interval that coincides with the length of a turn of the DNA helix. Pederson et al (2014) attributed the 10 bp read periodicity in specimens greater than 4,000 years old to protection of the DNA by nucleosomes preferentially positioned at 10 bp intervals. We observed a striking periodicity pattern averaging 10.8 bp in HA extracted samples and 10.1bp in the proK-PC samples. The shorter periodicity in the proK treated samples may be due to reduced protection of the ends of DNA fragments by digestion of the nucleosomes during extraction. We did not observe a signal of nucleosome occupancy in read depth or in enrichment of fragments of nucleosome length (147 bp) as did Pederson et al., perhaps because we sequenced shorter fragments to comparatively low depth. However, the appearance of 10 bp periodicity suggests it may be possible to infer nucleosome occupancy from patterns of DNA degradation observed in formalin-preserved specimens if higher coverage is achieved.

**Figure 5.**
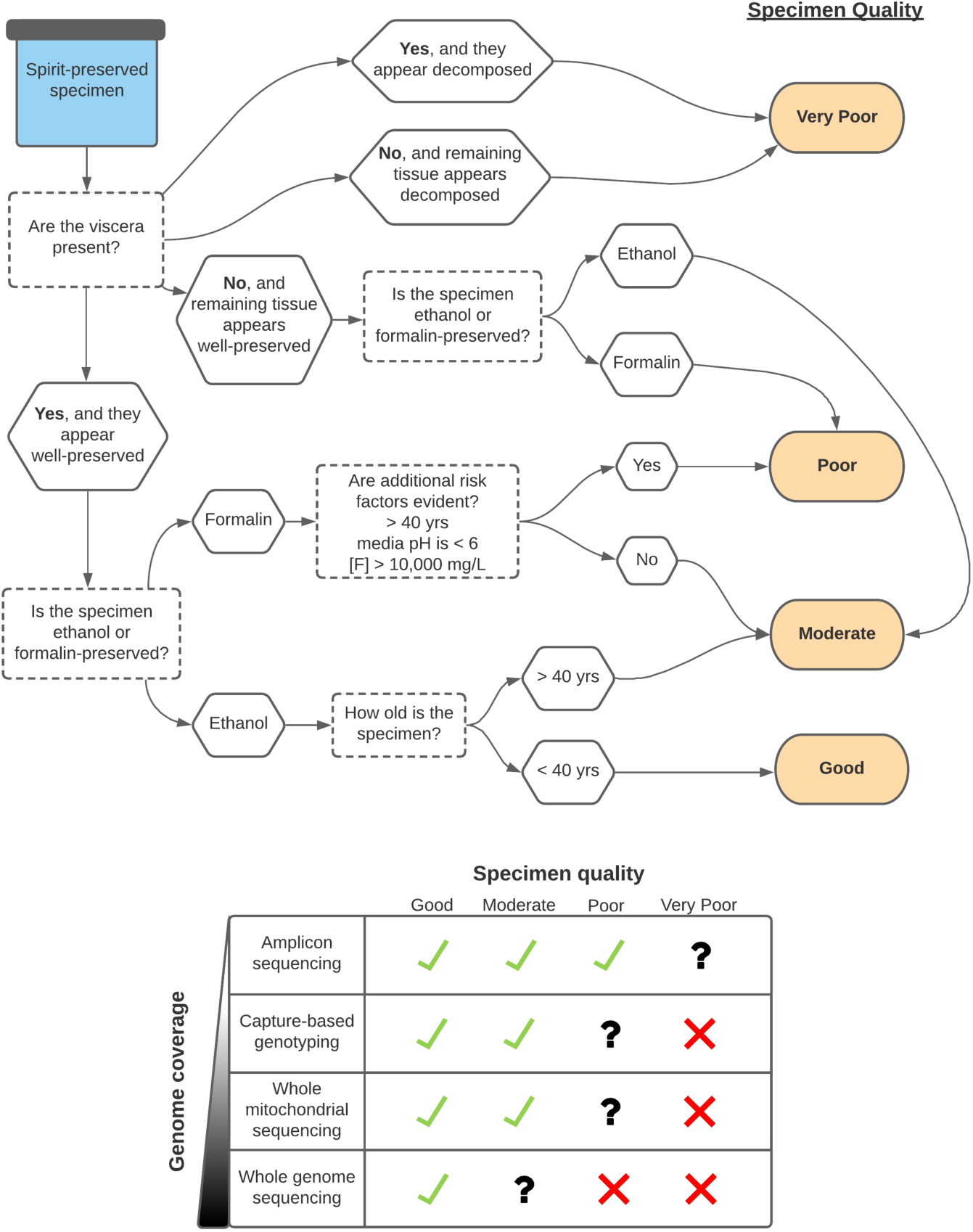
Decision-making tree for a priori estimation of likely sequencing success in spirit-preserved museum specimens. Green ticks indicate the specimen is well-suited the sequencing application and there is a high likelihood of success. Black question marks indicate the specimen is marginal for the sequencing application and there is high variation in the likelihood of success. Red crosses indicate the specimen is not well-suited for the sequencing application and there is a low likelihood of success.

### Managing expectations

We have shown WGS of formalin-preserved museum specimens is feasible and success can be improved through specimen quality vetting. We stress; however, measures of specimen quality are imperfect and the key parameters may vary between and within museum collections. Modern collection institutions aim to limit light exposure and temperature variation within their spirit vaults. With older specimens, the likelihood they have been exposed to undocumented DNA-degrading conditions increases. We found the age of the specimen was not strongly predictive of sequencing success, however, we did not sample specimens collected prior to the 1960s. This warrants further investigation into the extent to which intact DNA can be extracted from much older formalin-preserved specimens.

While preservation media pH and [F] were not predictive of sequencing success in our specimens, we note these measures do not always accurately reflect preservation condition. Most institutions periodically top up the specimen jars in their spirit vaults to replace ethanol lost through evaporation. In some cases, the preservation media is replaced entirely. Thus, media pH and [F] values at the time of sampling for sequencing may not reflect preservation and long-term storage conditions. With additional sampling of older and more varied specimens, it may be possible to establish clear correlates of sequencing success associated with pH and [F].

Both researchers and museums would benefit from an improved set of guidelines for strategic decision making based on independent quality metrics rather than qualitative *ad hoc* assessments. This will empower researchers to most effectively deploy their sequencing budgets and support museums in deciding when to grant requests for destructive sampling. A cost-benefit analysis should be conducted prior to genomic sequencing of museum specimens. From the perspective of the museum, destructive sampling should be avoided if the specimen is unlikely to yield sufficient DNA to achieve a project’s aims. From the perspective of the researcher, sequencing of high-quality specimens should be prioritized to generate high-quality data. To assist in making these assessments, we provide a decision-making tree (Figure 5) for use by both curators and researchers to determine which specimens are likely to be appropriate for genomic analyses.

Ultimately, museum curators decide if the potential benefit of sequencing outweighs the damage to the specimen through destructive sampling. Once sampling and DNA extraction has been completed, the decision to proceed with library preparation and sequencing can be made on the basis of DNA yield. We found specimens with high DNA yield (> 1,500 ng/50 mg tissue) produced a high percentage (> 20%) of mappable reads while specimens with low DNA yield (< 200 ng/50 mg tissue) produced virtually no mappable reads. While specimens yielding between 200–1,500 ng of DNA per 50 mg tissue produced relatively low genomic coverage, they did produce high coverage of the mitochondrial genome. Thus, reconstruction of historical mitochondrial haplotypes may be possible from specimens yielding low quantities of DNA. When nuclear data is required, high-volume sequencing should be reserved for high-quality specimens. Generally speaking, most research projects aim to sequence a small number of museum specimens with which to provide a base-line for comparison to contemporary specimens. In light of the limited availability of historical specimens in collections, it is often reasonable and feasible to allocate a relatively large budget to conduct deep sequencing of a small number of specimens.

## Conclusions

Our results demonstrate formalin-fixation is not a complete barrier to WGS in museum specimens. While success is not a guarantee, the use of HA lysis for DNA extraction followed by an appropriate sequencing library preparation optimized for degraded DNA can produce libraries of sufficient complexity for genomic analyses. When selecting specimens for sequencing, our results indicate those with poor gut integrity are least likely to yield sufficient DNA for sequencing.

## Methods

### Preservation media condition survey

We conducted an unbiased survey of the ANWC spirit vault to measure variation in preservation characteristics that can be sampled without disturbing the specimen. We randomly selected 149 specimen jars spanning a range of taxonomic groups and ages, and removed a 25 mL aliquot of preservation media. We measured pH using an Orion^TM^ Versa Star Pro^TM^ benchtop pH meter (*Thermo Scientific*) and [F] using MQuant® test strips (*Merck*). Where [F] was at the upper detection limit of the test strips, we diluted the aliquot 1:10 with ultrapure water and remeasured, extrapolating the neat concentration of the media by multiplying the measurement by the dilution factor.

### Specimen selection

To select specimens for genomic sequencing, we first identified those with a publicly available whole-genome reference for the specimen species or closely related species. Of these specimens, we selected 12 representing a range of taxonomic groups, preservation conditions and ages and sampled 50 mg of tissue. We sampled liver tissue when it was available. Muscle was sampled from an ethanol-preserved *P. cinereus* specimen and from formalin-preserved *T. guttata* and *O. anatinus* specimens. Skin was sampled from an ethanol-preserved *P. carbo*. All specimens sampled with liver were preserved as whole animals whereas substantial portions of the body were absent from those specimens sampled with muscle or skin (Supplementary Figure 2). From the nine formalin-preserved specimens, we selected three with which to test the relative success of three DNA extraction methods. To represent “good” quality formalin-preserved specimens, we selected a cane toad (*R. marina*) preserved in 2002. Visually, this specimen appeared minimally degraded and measurements of the storage media indicated low [F] and a neutral pH. To represent “poor” quality formalin-preserved specimens, we selected a tammar wallaby (*M. eugenii*) preserved in 1989 and a saltwater crocodile (*C. porosus*) preserved in 1973. Visually, these two “poor” specimens were reasonably well-preserved, however, measurements of the storage media indicated substantial [F] in both specimen jars and mildly acidic pH in that of the wallaby.

### Tissue preparation

Prior to DNA extraction, we liquid nitrogen pulverized all dissected tissue into a fine powder using a cryoPREP® (*Covaris*) dry pulverizer (three impacts to a TT05 tissueTUBE^TM^ on intensity setting three; 10 sec in liquid nitrogen between impacts). We then stored the pulverized tissue powder in 70% ethanol at -80°C until further processing. We re-hydrated the pulverized tissue by stepping it into 50% ethanol, 30% ethanol then TE buffer with rocking for 10 min intervals. For the nine formalin-fixed tissues, we quenched excess formaldehyde by rocking for 2 hrs in 1 mL GTE buffer (100 mM glycine, 10 mM Tris-HCL, pH 8.0, 1 mM EDTA), followed by a further wash in fresh GTE for 2 hrs and a final fresh GTE wash overnight at room temperature. We removed the GTE buffer and washed with rocking in sterile water for 10 min.

### Proteinase K DNA extraction

We conducted two variations on a standard proteinase K (proK) digestion. For each specimen, we digested two 50 mg (wet weight) aliquots of tissue overnight at 55°C with 30 µL of 20 mg/mL proteinase K in 970 µL lysis buffer (10 mM NaCl, 20 mM Tris-HCl, pH 8.0, 1 mM EDTA, 1% SDS). We isolated DNA from the proK lysates with either (A) three extractions of phenol-chloroform followed by ethanol precipitation (proK-PC), resuspending the DNA in 30 µL TE, or (B) a QIAquick PCR purification column (*Qiagen*) (proK-col), following the manufacturer’s instructions and eluting the DNA in 30 µL TE. Alongside the museum tissues, we processed tissue-free controls. We quantified extracted dsDNA using a Qubit fluorometer and high sensitivity (HS) DNA kit (*Invitrogen*).

### Hot alkaline lysis DNA extraction

For the hot alkaline lysis (HA) extractions, we heated 50 mg (wet weight) tissue aliquots to 120°C for 25 min in 500 µL of alkali buffer (0.1 M NaOH with 1% SDS, pH 13) according to methods described in (52). We purified DNA from the lysate with three phenol-chloroform extractions followed by ethanol precipitation, resuspending the DNA in 30 µL TE. Alongside the museum tissues, we processed tissue-free controls. We quantified extracted dsDNA using a Qubit fluorometer and HS DNA kit.

### Library preparation methods

To avoid cross-contamination, we prepared all sequencing libraries in the Ecogenomics and Bioinformatics Laboratory trace facility at the Australian National University following standard anti-contamination procedures. We prepared libraries from all DNA extracts and tissue-free controls using two methods developed for high efficiency conversion of fragmented aDNA; the single-stranded method v2.0 (ss2) (25) and the BEST double-stranded method (dsBEST) (26). Concurrently, we prepared DNA-free control libraries. For sequencing of Read 1 in both library preparation methods, we used an adapter with the sequence 5’– AGATCGGAAGAGCACACGTCTGAACTCCAGTCAC–3’. For sequencing of Read 2, we used adapters with the sequences 5’–GGAAGAGCGTCGTGTAGGGAAAGAGTGT–3’ and 5’–AGATCGGAAGAGCGTCGTGTAGGGAAAGAGTGT–3’ for the ss2 and dsBEST methods, respectively. We removed excess adapter and primer dimer by isolating fragments between 160 bp and 400 bp from the resulting libraries using the PippinHT size-selection system (*Sage Science*). We further purified the libraries with a MinElute PCR purification kit (*Qiagen)* and quantitated the library concentrations using the LabChip GXII (*PerkinElmer*) capillary electrophoresis system. We then pooled the libraries in approximately equimolar concentrations and measured the concentration of the final pooled library using a Qubit fluorometer and HS DNA kit. The Australian Genome Research Facility sequenced the pooled library on a 150 bp paired-end S4 flow cell on the Illumina NovaSeq 6000 platform.

### Quality control of raw reads

We computed quality control metrics for the raw reads using FastQC v.0.11.8 (60). Our adapter content analysis included both default Illumina adapters and our custom library adapters. To rapidly detect library contamination by non-target species’ DNA, we classified the taxonomic origin of reads using Kraken2 v.2.0.9b (61). We estimated the number of unique fragments present in the raw sequence libraries with the EstimateLibraryComplexity function of PICARD v.2.9.2 (62).

### Alignment

We aligned reads to reference nuclear and mitochondrial genomes obtained from the DNA Zoo Consortium (63, 64) and GenBank (65) (Supplementary Table 1). Species-specific reference genomes were not available for three of the specimens. For *A. audax*, *F. cenchroides* and *P. minima,* we used the reference genomes of species in the same genera- *A. chrysaetos*, *F. perigrinus* and *P. vitticeps*, respectively (Supplementary Table 1). We hard-masked the eleven genomes with RepeatMasker v.4.1.0 (66) including our ss2 and dsBEST library adapters in the repeat database and applying the -qq option allowing 10% less sensitivity while decreasing processing time. We aligned raw reads with the kalign function of the ngskit4b tool suite v.200218 (67) with options -c25 (--minchimeric= <INT>; minimum chimeric length as a percentage of probe length) -l25 (--minacceptreadlen= <INT>; after any end trimming only accept read for further processing if read is at least this length) -d50 (--pairminlen= <INT>; accept paired end alignments with observed insert sizes of at least this) -U4 (--pemode= <INT>; paired end processing mode: 4 - paired end no orphan recovery treating orphan ends as SE). We removed PCR and optical duplicates from the alignments using the MarkDuplicates function of PICARD enabling REMOVE_DUPLICATES=TRUE. For each de-duplicated alignment, we computed a histogram of aligned insert lengths and calculated the mean aligned insert length using the CollectInsertSizeMetrics function of PICARD.

### Genome coverage analyses

We estimated nuclear genome coverage (C_nuc_) as the number of unique aligned reads multiplied by the mean insert length divided by unmasked genome size. To estimate how much genomic coverage could be achieved by increasing sequencing depth, we calculated the sequenced proportion of the prepared library as the number of read pairs examined divided by the estimated library size. We estimated the number of possible reads represented in the prepared library by dividing the number of actual reads aligned by the sequenced proportion of the library. We then roughly estimated the potential genomic coverage represented in the full prepared library (C_pot_) as: (# possible reads x mean insert length (bp)) ÷ genome size (bp). To calculate the proportion of mitochondrial genome sites with 30X or greater coverage (C_mt_), we executed the Samtools *depth* function (68) on SAM files for the mitochondrial contigs for each species combined across all libraries.

### Statistical analyses

We performed statistical analyses in the R environment, v.4.0.2 (69) and produced figures using the packages *ggplot2* (70) and *ggpubr* (71). To test if the residuals of data were normally distributed, we ran Shapiro-Wilk tests with the function *shapiro.test*. We conducted T-tests with the function *t.test*, analyses of variance (ANOVA) with the function *aov* and computed confidence intervals using Tukey’s Honest Significant Difference method (Tukey test) with the function *TukeyHSD* in the base package *stats*. We computed Pearson correlation coefficients with associated p-values with the *ggpubr* function *stat_cor*.

## Declarations

### Ethics approval and consent to participate

Not applicable.

### Consent for publication

Not applicable.

### Availability of data and materials

The sequencing data generated and analysed in this study are archived in the <u>CSIRO Data</u> <u>Access Portal</u>. Correspondence and requests for materials should be addressed to CEH

## Funding

Funding for this study was provided by the Environomics CSIRO Future Science Platform (grants R-10011 and R-14486) awarded to CEH.

### Authors’ contributions

This study was conceived by CEH. Experiments were designed by CEH, MRA and AG and conducted by MRA and AG. Data analysis was conducted by EEH, JS, MRA and AG and advised by DMG and CEH. All authors contributed to the writing and editing of the manuscript.

## Acknowledgements

We thank Olly Berry and Andrew Young for their leadership within the Environomics Future Science Platform. We thank the director of the <u>Australian National Wildlife Collection</u>, Leo Joseph, and the ANWC staff (specifically, Margaret Cawsey, Alex Drew, Tonya Haff, Dave Spratt and Chris Wilson) for their contributions of curatorial expertise, metadata management and sampling assistance. We thank Kerensa McElroy for her assistance and guidance in data management. We thank Ondrej Hlinka and CSIRO IM&T Client Services for their assistance in utilising the CSIRO Pearcey supercomputing system. We thank Niccy Aitkin for her guidance in utilising the Australian National University’s Ecogenomics and Bioinformatics Laboratory for library preparation. We thank the Australian Genome Research Facility for their conversations around sequencing. We thank Sharon Appleyard, Meghan Castelli, Andrew George, Peter Grewe, Michael Hope, Safia Maher, Annette McGrath, Corinna Paeper, Cheng Soon-Ong, Andrew Spriggs, Jen Taylor and Christfried Webers for their valuable comments on the study design and implementation. We would like to acknowledge the contribution of Bioplatforms Australia in the generation of data used in this publication. Bioplatforms Australia is enabled by NCRIS.

