## Supplemental Figure 1 for "Unlocking inaccessible historical genomes preserved in formalin"

### Supplementary Figures

#### Supplementary Figure 1.  Collection survey summary

Range of pH (A) and age (B) of ethanol- and formalin-preserved specimens; Outliers are indicated with diamonds, **** = $p < 0.0001$

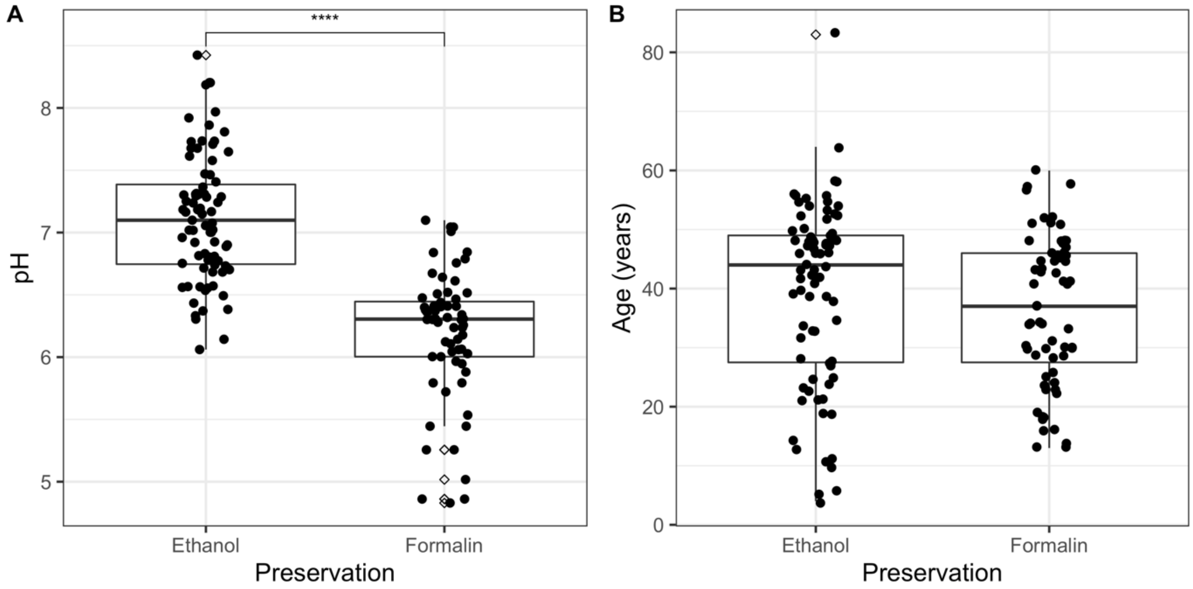
