## Supplementary figures and images for "Unlocking inaccessible historical genomes preserved in formalin"

### Supplemental Figure 2

#### Supplementary Figure 2. Specimen photographs

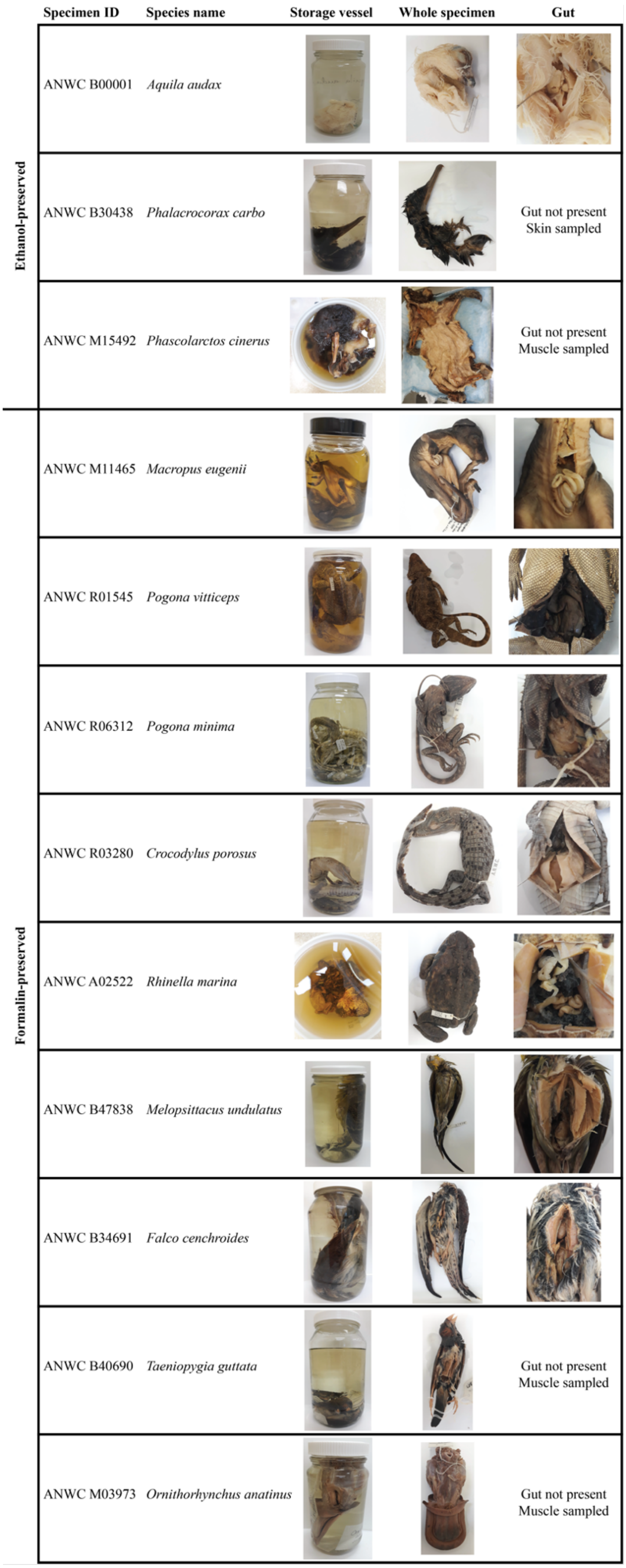
