## Supplemental Figure 3 for "Unlocking inaccessible historical genomes preserved in formalin"

**Supplementary Figure 3. DNA yield summary**

DNA yield expressed as a function of (A) extraction method as tested on the *R. marina*, *M. eugenii* and *C. porosus* specimens and (B) preservation method across all 12 specimens. DNA yield from the formalin-preserved specimens as expressed as a function of (C) pH, (D) tissue type ($p < 0.05$) (E) formaldehyde concentration and (F) age. Outliers are indicated with asterisks in B. In E and F, R = Pearson's correlation coefficient.

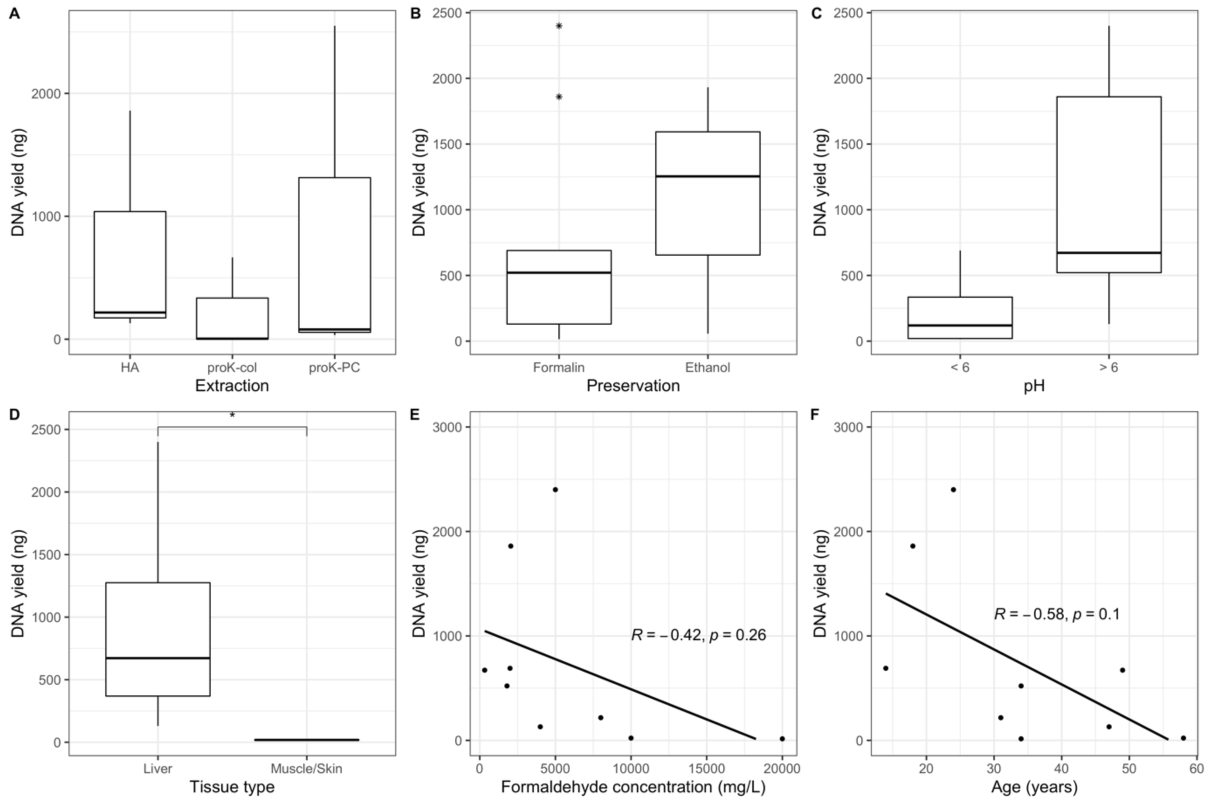
