## Supplemental Table 1 for "Unlocking inaccessible historical genomes preserved in formalin"

**Supplementary Table 1. Reference genomes used in this study**

Genome sequences were sourced from GenBank unless noted (ϕ), where genomes were sourced from DNA Zoo (https://www.dnazoo.org).

| Species name | Species-specific genome? | Accession/version | Mitochondrial genome accession | Nuclear genome size (Gb) | Mitochondrial genome size (Kb) |
|---|---|---|---|---|---|
| *Aquila audax* | No, used *Aquila chrysaetos* | Aquila_chrysaetos-1.0.2_HiC ϕ | NC_024087.1 | 1.3 | 17.3 |
| *Crocodylus porosus* | Yes | GCA_001723895.1 | NC_008143.1 | 2.8 | 17.3 |
| *Falco cenchroides* | No, used *Falco peregrinus* | GCA_001887755.1 | NC_000878.1 | 1.2 | 18.1 |
| *Macropus eugenii* | Yes | me-1k ϕ | KJ868119.1 | 2.9 | 16.9 |
| *Melopsittacus undulatus* | Yes | GCA_000238935.1 | NC_009134.1 | 1.2 | 18.2 |
| *Ornithorhynchus anatinus* | Yes | GCA_004115215.1 | NC_000891.1 | 2.3 | 17.0 |
| *Phalacrocorax carbo* | Yes | GCA_000708925.1 | NC_027267.1 | 1.1 | 19.0 |
| *Phascolarctos cinereus* | Yes | GCA_002099425.1 | NC_008133.1 | 3.5 | 16.4 |
| *Pogona minima* | No, used *Pogona vitticeps* | GCA_900067755.1 | NC_006922.1 | 1.8 | 16.8 |
| *Pogona vitticeps* | Yes | GCA_900067755.1 | NC_006922.1 | 1.8 | 16.8 |
| *Rhinella marina* | Yes | GCA_900303285.1 | Pers. Comm. Rollins 2021 | 2.6 | 18.2 |
| *Taeniopygia guttata* | Yes | GCA_003957565.1 | NC_007897.1 | 1.2 | 16.9 |
