## Supplemental Table 2 for "Unlocking inaccessible historical genomes preserved in formalin"

### Supplementary Table 2. Kraken results

| Species name | Extraction method | Library prep | Proportion of reads with matches to taxon group: Genus *Mus* | Other notable hits |
|---|---|---|---|---|
| Negative control | HA | dsBEST | 0.3 | 1.8% *Aquila chrysaetos* |
| Negative control | proK-col | dsBEST | 19.4 | 6.5% *Aquila chrysaetos* |
| Negative control | proK-PC | dsBEST | 5.9 | 2.1% *Aquila chrysaetos* |
| Negative control | HA | ss2 | 7.3 | 3.3% *Homo sapiens*; 3.3% *Canis lupus* |
| Negative control | proK-col | ss2 | 0.9 | |
| Negative control | proK-PC | ss2 | 15.9 | 1.5% *Homo sapiens* |
| *Rhinella marinus* | HA | dsBEST | 0.1 | |
| *Rhinella marinus* | proK-col | dsBEST | 0.1 | |
| *Rhinella marinus* | proK-PC | dsBEST | 0.6 | |
| *Rhinella marinus* | HA | ss2 | 0.1 | |
| *Rhinella marinus* | proK-col | ss2 | 0.1 | |
| *Rhinella marinus* | proK-PC | ss2 | 0.1 | |
| *Macropus eugenii* | HA | dsBEST | 0.1 | |
| *Macropus eugenii* | proK-col | dsBEST | 0.8 | |
| *Macropus eugenii* | proK-PC | dsBEST | 2.1 | |
| *Macropus eugenii* | HA | ss2 | 0.1 | |
| *Macropus eugenii* | proK-col | ss2 | 2.7 | 4.6% *Homo sapiens*; 0.5% *Aquila chrysaetos* |
| *Macropus eugenii* | proK-PC | ss2 | 0.1 | |
| *Crocodylus porosus* | HA | dsBEST | 2.3 | |
| *Crocodylus porosus* | proK-col | dsBEST | 75.6 | 1.4% *Homo sapiens* |
| *Crocodylus porosus* | proK-PC | dsBEST | 22.1 | |
| *Crocodylus porosus* | HA | ss2 | 10.4 | 2.8% *Aquila chrysaetos* |
| *Crocodylus porosus* | proK-col | ss2 | 58.2 | 2.1% *Homo sapiens* |
| *Crocodylus porosus* | proK-PC | ss2 | 2.6 | |
| *Melopsittacus undulatus* | HA | dsBEST | 0.8 | 2.0% *Aquila chrysaetos* |
| *Melopsittacus undulatus* | HA | ss2 | 0.4 | |
| *Pogona minima* | HA | dsBEST | 0.1 | |
| *Pogona minima* | HA | ss2 | 0.5 | |
| *Pogona vitticeps* | HA | dsBEST | 0.2 | |
| *Pogona vitticeps* | HA | ss2 | 0.1 | |
| *Phalacrocorax carbo* | HA | dsBEST | 0.4 | |
| *Phalacrocorax carbo* | HA | ss2 | 0.1 | |
| *Taeniopygia guttata* | HA | dsBEST | 0 | |
| *Taeniopygia guttata* | HA | ss2 | 0.3 | |
| *Aquila audax* | HA | dsBEST | 0.2 | |
| *Aquila audax* | HA | ss2 | 0.3 | |
| *Falco cenchroides* | HA | dsBEST | 0.1 | |
| *Falco cenchroides* | HA | ss2 | 0.3 | |
| *Phascolarctos cinereus* | HA | dsBEST | 0.2 | |
| *Phascolarctos cinereus* | HA | ss2 | 0.3 | |
| *Ornithorhynchus anatinus* | HA | dsBEST | 14.4 | 9.7% *Homo sapiens* |
| *Ornithorhynchus anatinus* | HA | ss2 | 9.5 | 25.1% *Homo sapiens* |
